# Dissecting the genetic architecture of sunflower head diameter using genome-wide association study

**DOI:** 10.1101/2022.10.24.513623

**Authors:** Yavuz Delen, Ravi V. Mural, Gen Xu, Semra Palali Delen, James C. Schnable, Jinliang Yang, Ismail Dweikat

**Author notes:** Corresponding authors: Jinliang Yang and Ismail Dweikat.

## Abstract

Sunflower (*Helianthus annuus* L.) plays an essential role in meeting the edible oil demand worldwide. Sunflower seed yield can be decomposed into several yield component traits, one of which is the head diameter. In 2019, 2020, and 2022, we evaluated the head diameter phenotypic variation on a set of diverse sunflower accessions (N=342) in replicated field trials. By combining three years of field data, the broad sense heritability (*H*^2^) of the head diameter trait was estimated to be 0.88. Then, a subset of N=274 accessions was genotyped by using the tunable genotyping-by-sequencing (tGBS) method, resulting in 226,779 high-quality SNPs. With these SNPs and the head diameter phenotype, the genome-wide association study (GWAS) was conducted using two statistical approaches: 1) the mixed linear model (MLM) and 2) the fixed and random model circulating probability unification (farmCPU). The MLM and farmCPU GWAS approaches identified 106 and 8 significant SNPs that were placed close to 53 and 21 genes, respectively. Two significant peaks were identified in MLM, with a strong signal on chromosome 10 and a less strong signal on chromosome 16. The farmCPU method detected the same signals on chromosomes 10 and 16 and several additional significant signals on other chromosomes. The head diameter associated genetic loci and the underlying candidate genes can be leveraged for further functional validation and serve as a basis for sunflower oil yield improvement.

## Introduction

Sunflower (*Helianthus annuus* L.), a member of the *Helianthus* genus from the Compositae (Asteraceae) family, is a diploid crop with 2n=34 chromosomes. It is native to North America and was initially domesticated by Native Americans in the East Central region of the US (1). Sunflower is the second-largest hybrid crop after maize (*Zea mays* L.) (2) and fourth largest crop in oil production after palm, soybean, and rapeseed (3), providing 25-50% edible oil in seeds (4). Currently, the majority of sunflower hybrids contain 45–50 % oil in their seeds (5). The ability of sunflowers to adapt to a wide range of environments has resulted in its cultivation around the globe on every continent except for Antarctica (2). It is now cultivated in more than 70 countries across the world (6), providing more than 13% of the total edible oil globally (7). A wide range of phenotypic variation in terms of the desired traits (i.e., seed size, head diameter, oil content, resistance to biotic and abiotic factors) and the genetic diversity in the domesticated and wild sunflower populations provide a valuable source for sunflower breeding programs and genetic studies. Sunflower’s ability to grow in disrupted and marginal agricultural land suggests it may play an increasingly crucial role in meeting the needs of increasing world population.

Sunflowers are mostly grown for seed production to extract oil from the seeds or for confectionery purposes. Even though the confectionery-type seeds involve higher protein than the seeds used for oil production, they have a low amount of oil than the oil types (8). Therefore, the seeds used in oil production are called oilseeds, and the confectionery-type seeds are named non-oil seeds. The sunflower seeds used for oil production are generally smaller than those for confectionery types. The sunflower seed yield is affected by various traits like seed size and seed weight. In addition to those, floral traits play an essential role in seed yield. They influence pollination success and eventually, sunflower seed yield (9; 10). The sunflower blossom is called the sunflower head (Capitulum). Wild progenitors of sunflower are highly branched and produce many heads on the branches while domesticated sunflower is unbranched and produces mostly single head. Each sunflower head includes disk flowers and a line of ray flowers on the outer side of the disk flowers. The sunflower heads consist of up to 3,000 tubular disk flowers, which are fertile and produce the fruit known as cypsela or achene (8). The non-oilseed hybrids with up to 8,000 disk flowers are also available (11). In contrast to the disk flowers, the ray flowers that are called petals do not bear anthers, stigmas, or ovules. Therefore, they are sterile and do not produce any seed, however, play an essential role in attracting pollinators (12–14) as an evolutionary trade-off to increase pollination and seed production (15).

As a floral trait, the head diameter trait of the sunflower is one of the most important traits since it impacts sunflower seed yield. It has been indicated that head size and the number of seeds are highly correlated with each other (16–18), contributing directly sunflower seed yield (19; 20). High variation in head size and head shape (flat, concave, convex, etc.) presents in the sunflower populations. In contrast to wild progenitors, domesticated sunflowers have wider heads (21–23). Studies show that sunflower head diameter is a quantitative trait affected by many genetic loci, including both additive and non-additive genetic components (24–26). To assess the genetic mechanism of sunflower head diameter, many quantitative trait loci (QTL) studies have been conducted to identify loci regulating this trait. However, genome-wide association studies (GWAS) for sunflower head diameter have been comparatively less common than QTL studies. Sunflower has a relatively big genome size (3.6 Gb), and its genome assembly that is used as a reference genome was just released in 2017 (27). With the technological advances, the cost effective and easily reachable genotyping and sequencing opportunities have opened new perspectives to plant breeding and genetics programs by enabling genome-wide studies. Recently, a number of GWAS have been conducted on sunflower traits (27–36), resulting in hundreds of trait-associated loci. Among those studies, only Dowell et al. (2019) (32) focused on the head diameter trait and used about 6,000 markers to detect associations with the head (disk) size by GWAS. Their study found trait-associated loci in large genomic regions. Higher resolution GWAS, however, is needed to further dissect the genetic components in determining the head diameter traits.

This study aims to reveal the genetic basis controlling sunflower head diameter variation using a larger set of SNP markers. For this purpose, we genotyped a sunflower diversity panel using tGBS methods and obtained 226,779 high quality SNPs. Meanwhile, we collected the phenotypic data of this diversity panel in the years of 2019, 2020, and 2022 in the replicated field trials. Then, we conducted GWAS and identified 106 and 8 significant SNPs using the MLM and farmCPU methods, which were located close to 53 and 21 genes, respectively. These trait-associated SNPs and the candidate genes underneath the GWAS peaks can be potentially applied for further sunflower improvement.

## Materials and methods

### Plant materials and field experimental design

A set of 342 sunflower accessions, originally collected in 25 different countries on six continents, were obtained from the North Central Regional Introduction Station (NCRPIS) in Ames, Iowa, USA. Accessions originated from diverse geographical locations were randomly selected to maximize the genetic diversity.

These accessions were grown at the University of Nebraska-Lincoln’s research farm at Havelock in 2019 (40^◦^ 51^′^ 15.9^′′^ *N*, 96^◦^ 36^′^ 42.6^′′^ *W*), 2020 (40^◦^ 51^′^ 26.5^′′^ *N*, 96^◦^ 36^′^ 53.4^′′^ *W*), and 2022 (40^◦^ 51^′^ 20.1^′′^ *N*, 96^◦^ 36^′^ 32.4^′′^ *W*). For the field experiment, an incomplete block design was employed with two main blocks, four split plots per block, and three replicates of the check (PI 432513) per split plot. Each genotype was planted in a 3.6 meter long single row with 0.75 meter row spacing and an alleyway of 0.9 meters. Twelve seeds were planted per row, leading to a density of about 45,000 plants per hectare. Head diameter (in centimeters) was measured for three representative plants per plot, excluding edge plants. Measurements were carried out after pollination but prior to maturity.

### Phenotypic data processing and heritability calculation

Best Linear Unbiased Prediction (BLUP) calculation for head diameter was calculated using lme4 package (37) in R (v 4.2.0) (38). In the analysis, the following model was fit using phenotype data collected in 2019, 2020, and 2022: *Y ∼* (1 | *genotype*) + (1 | *block*) + (1 | *split − plot*) + (1 | *year*) + (1 | *genotype* : *year*) + *error*, where *Y* represents the phenotype (Sunflower head diameter). In this study, genotype, block, split-plot, year, and genotype-by-year interaction were treated as random effects. The BLUP calculated using the combination of data collected in 2019, 2020, and 2022 was used in the GWAS analysis.

In the BLUP model,

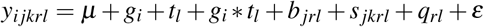

where *y*_*i jkrl*_ refers the phenotypic value of the *i*^*th*^ genotype evaluated in the *k*^*th*^ split-plot of the *j*^*th*^ block of *r*^*th*^ replicate nested within the *l*^*th*^ year; *µ* is the overall mean; *g*_*i*_ is the random effect of the *i*^*th*^ genotype; *t*_*l*_ is the random effect of the *l*^*th*^ year; *g*_*i*_ *t*_*l*_ is the random effect of the *i*^*th*^ genotype with the *l*^*th*^ year interaction; *b* _*jrl*_ is the random effect of the *j*^*th*^ block of the *r*^*th*^ replicate within the *l*^*th*^ year; *s* _*jkrl*_ is the random effect of the *k*^*th*^ split-plot of the *j*^*th*^ block of the *r*^*th*^ replicate within the *l*^*th*^ year; *q*_*rl*_ is the random effect of the *r*^*th*^ replicate nested within the *l*^*th*^ year; *ε* is the random residual error.

Broad-sense heritability (*H*^2^) was calculated based on the equation *H*^2^ = *V*_*G*_*/V*_*P*_ (39; 40), where *V*_*P*_ is *V*_*G*_+*V*_*E*_, *V*_*G*_ is total genetic variance, *V*_*P*_ is total phenotypic variance, and *V*_*E*_ is phenotypic variance due to environmental factors. Regarding this, the broad-sense heritability of sunflower head size was calculated for combined environments of 2019, 2020, and 2022 by the following equation.

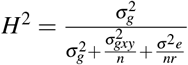

where 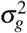 is the components of variance for genotype, 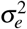 is the components of variance for environment, 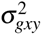 is the components of variance for genotype by year interaction, *n* is the number of years, and *r* is replications.

### DNA extraction and genotyping

All accessions were grown in the greenhouse to collect leaf samples for DNA extraction. The accessions were grown on 10×10 cm pots using standard greenhouse mix with a ratio of 5 gallons of peat, 3 gallons of soil, 2.5 gallons of sand, and 2.5 gallons of vermiculite. Two weeks after planting, samples (600-700 mg) from young leaves at the V4 stage were collected in sterile Eppendorf tubes, and then samples were placed in a − 80^◦^ Celsius freezer. Frozen samples were lyophilized using a freeze-drying machine (VirTis 25XL) for three days at − 70^◦^ Celsius. The lyophilized tissue samples/leaf samples were transferred into the tubes of three 96-well collection kits and shipped to Freedom Markers at Ames, Iowa (https://www.freedommarkers.com/) for genotyping. The accession “PI 490282” was placed once in each well plate as a check that can be used to confirm the quality of genotyping. The company used the BioSprint MagAttract 96 DNA Plant Core Kit by QIAGEN for DNA extraction and utilized a PicoGreen kit on an Eppendorf Plate Reader for assessing purity and quality control. The SNP genotyping was conducted by Freedom Markers using tGBS^®^ genotyping by sequencing technology utilizing restriction enzyme Bsp1286I (41). The tGBS libraries were constructed using the QCd DNA (as described above) from each sample and the resulting libraries were subsequently sequenced on an Illumina HiSeq X with 2 × 150 bp paired-end reads. Due to the high missing rate, 11 of the 285 samples (PI_219650, PI_262519, PI_262521, PI_432522, PI_531502, PI_546360, PI_552946, PI_599770, PI_599781, PI_632338, and PI_650658) that were selected for genotyping were removed in subsequent analyses.

### Sequence data processing and SNP calling

The resulting fastq sequence data was scanned and quality-trimmed by Freedom Markers using an internal software that removes low-quality regions in a two-step process. In the first step, reads were scanned beginning at each end. If the nucleotide has PHRED quality values less than 15 (42; 43) they were discarded. In the second step, the read is scanned in 10 base pair windows and truncated if the average PHRED quality score of a 10 base pair window fell below the cutoff value of 15. These trimming parameters were adapted from trimming software “LUCY2” (44; 45).

Reads were aligned to the HanXRQr2.0-SUNRISE sunflower reference genome (https://www.ncbi.nlm.nih.gov/assembly/GCF_002127325.2/) (27) using GSNAP (46). Only confidently and uniquely mapped reads, defined as (≤ 2 mismatches every 36 bp and less than 5 bases for every 75 bp as tails) were used for subsequent analyses. SAMtools was used for conversion of alignment file formats (http://samtools.sourceforge.net/) (47).

Polymorphic sites that diverge from the reference genome in at least one sample were identified by utilizing all reads that uniquely align to the sunflower reference genome. After counting the reads at each SNP site that was considered to have potential, it was checked if the SNP site is interrogated meaning that a SNP site was supported by at least five unique reads. Freedom Markers reported the number of positions in the genome. If a segregating polymorphism exists at a certain location in the genome, a SNP marker was detected and genotyped.

A SNP was genotyped as homozygous in a given individual when at least 80% of all aligned reads at that site supported the most commonly observed nucleotide and at least five unique reads must support the most commonly observed nucleotide. For an individual to be genotyped as heterozygous at a given site at least 30% of all aligned reads that cover that position, and at least 5 unique reads must support each of the two most commonly observed nucleotides and at least 80% of all aligned reads covering that nucleotide site must support one of those two most commonly observed nucleotides. In both homozygous and heterozygous SNP criteria, polymorphisms in the first and last 3 bp of each read were not taken into consideration. In addition, both criteria include that at least a PHRED base quality value of 20, corresponding to an error rate of ≤ 1% must be estimated for the genotype call at that site in that individual.

To select the most proper set of SNPs for the imputation, several cut-offs were applied based on the minimum call rate (MCR). As a result, imputation was used on the MCR50 SNPs with 32.25% of missing data and 1.11% polymorphisms to fill in the gaps where there were insufficient reads to make genotype calls within a genotype. The maximum missing data rate per SNP site was 50% in the MCR50 SNPs (**Table 1**). Imputation was performed by using BeagleV5.1 (http://faculty.washington.edu/browning/beagle/beagle.html) with 50 phasing iterations and other default parameters. The following criteria in filtering MCR50 SNPs were applied: Minimum calling rate ≥ 50%, allele number = 2, number of genotypes ≥ 2, minor allele frequency (MAF) ≥ 5%, and heterozygosity rate range: 0% – (2 × *Frequency*_*allele*1_ × *Frequency*_*allele*2_ + 20%). To not cause the low accuracy associated with the imputation of scaffold SNPs, the imputation was performed only on the chromosome-based SNPs.

**Table 1.** Summary of SNPs genotyped in different MCR (Minimum Call Rate) levels.

|  | No. SNPs | Missing Data Points | % Polymorphisms<br>(Raw reads) |
| --- | --- | --- | --- |
| MCR50 | 247,008 | 21,986,571 / 68,174,208 = 32.25% | 247,008 / 22,154,507 = 1.11% |
| MCR60 | 169,294 | 12,300,433 / 46,725,144 = 26.33% | 169,294 / 14,595,270 = 1.16% |
| MCR70 | 100,121 | 5,601,229 / 27,633,396 = 20.27% | 100,121 / 8,493,851 = 1.18% |
| MCR80 | 43,928 | 1,702,588 / 12,124,128 = 14.04% | 43,928 / 3,835,705 = 1.15% |
| MCR90 | 8,533 | 166,432 / 2,355,108 = 7.07% | 8,533 / 886,584 = 0.96% |

### SNP analysis

Principal component analysis (PCA) was performed using TASSEL v5 (48). The eigenvalues of 10 PCs were used to create the scree plot to visualize the proportion of variance explained by each PC. The software PLINK 1.9 (49) was utilized in R (38) in order to estimate the linkage disequilibrium (LD) with the *r*^2^ statistics, build LD decay, and perform the minor allele frequency. In addition, the R package LDheatmap was used to plot the heat maps of pairwise LD between SNPs (50). Kinship, was estimated using TASSEL v5 (48) and a figure illustrating the kinship was created in Python (51). In addition, a neighbor-joining phylogenetic tree was constructed using the distance between each accession as per SNPs with MAF > 0.05 using a TASSEL plugin “-tree Neighbor” (TASSEL v5) (48). The phylogenetic tree was visualized using Interactive Tree Of Life (iTOL) (http://itol.embl.de/index.shtml), an online tool for displaying and annotating phylogenetic tree (52).

### Statistical methods for genome-wide association study

As a first step, 247,008 SNPs were detected and filtered into 246,671 SNPs by removing the excessive alignments on the unplaced genomic scaffolds. For the marker trait association analysis, the initial SNP set (246,671 SNPs) was filtered by removing the SNPs with minor allele frequencies of ≤ 0.05 across the 274 individuals, resulting in a set of 226,779 SNPs. The resulting marker set was employed for GWAS analysis by a single-locus model; MLM (P+K) (53) and a multi-locus model; farmCPU (54). Both models were run using the R package “rMVP” (v1.0.6) (55). For the MLM model, the kinship matrices having the relationship among individuals (56) and the first three principal components (PCs) were fit as covariates to control for the confounding effects of the population structure. The threshold for the significant association SNPs was set to 2.2×10^−7^(0.05/*n, n* = 226,779). For the farmCPU model, the kinship matrix calculated internally by the farmCPU algorithm was fitted as random effects in addition to the first three PCs as covariates. Manhattan plots and Q-Q plots representing GWAS results were plotted in rMVP itself.

## Results

### Phenotypic distribution, heritability, and BLUP value calculation

In this study, a set of 342 geographically widely distributed sunflower accessions were obtained from North Central Regional Introduction Station (NCRIS) (see **Supplementary Table S1**). Based on the location information extracted from the Germplasm Resources Information Network (GRIN), the majority 314/342 (92%) of the selected sunflower accessions originated from North America, Europe, and Asia, and a small subset 28/342 (8%) accessions from South America, Africa, Australia, or unknown locations (*n* = 2) (**Figure 1A**). These accessions were planted in 2019, 2020, and 2022 following an incomplete block design (see **Materials and Methods, Supplementary Figure S1**). From the field experiment, head diameter was manually measured from up to three representative plants per plot (**Supplementary Figure S2**).

**Figure 1.**
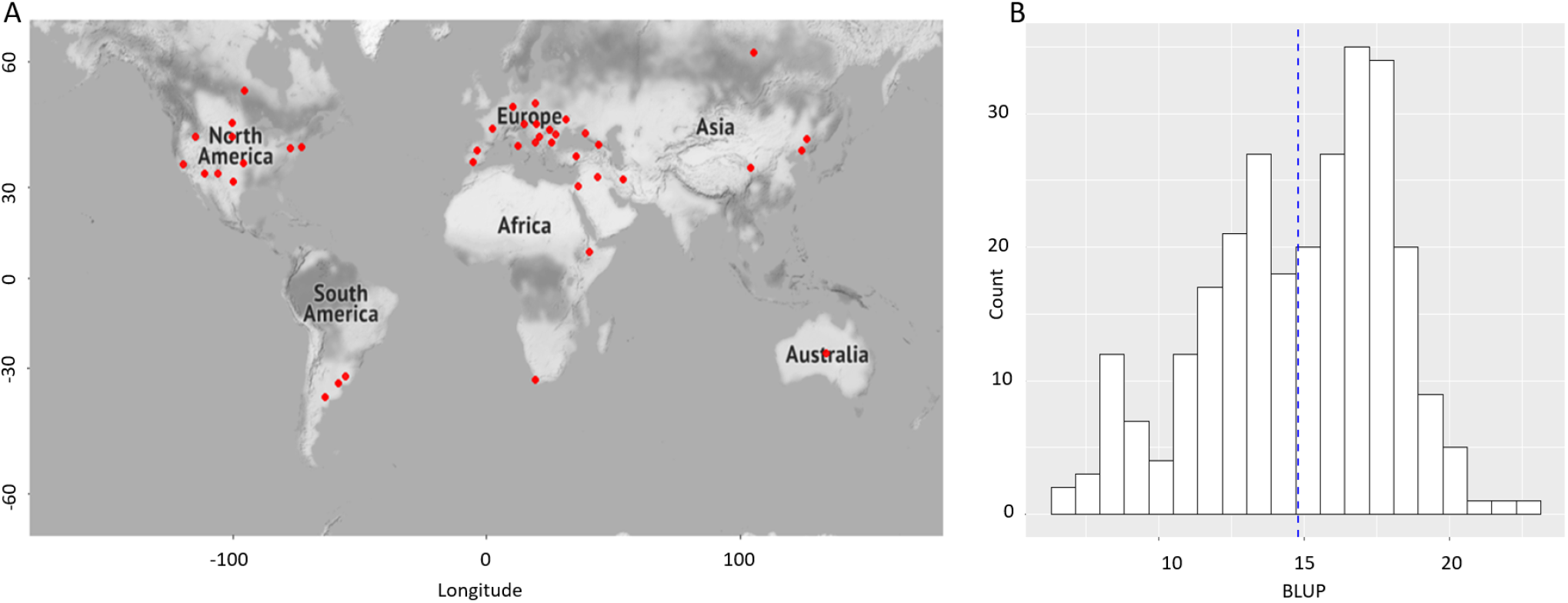
Geographic locations of genotyped sunflower accessions and the distribution of BLUP value. (**A**) A map illustrating the geographic coordinates of the sunflower accessions. Red dot denotes the location of each accession. (**B**) The histogram distribution of BLUP values. The blue dashed line shows the mean BLUP value of head diameter.

In 2019, as a result of more significant damage from sunflower moths, the distribution of observed sunflower head diameter was skewed towards substantially smaller heads (mean size ≈ 8 cm) as compared to 2020 (mean size ≈ 19 cm) and 2022 (mean size ≈ 17 cm) (**Supplementary Figure S3**). After analyzing these three years of data, the heritability (*H*^2^) of head diameter was estimated as 0.88. Excluding 2019 data obtained a slightly higher heritability of *H*^2^ = 0.92. Because the sunflower moth damage did not dramatically affect the heritability of the trait, we then combined three years of data to calculate BLUP values (**Materials and Methods**). As a result, BLUP values of the head diameter trait ranged from 6.8 to 22.8 with a mean of 14.8 (**Figure 1B**).

### SNP genotyping and population structure analysis

For a subset (285/342, or 83%) of the sunflower accessions, we conducted genotyping using the tGBS method (57) (**Materials and Methods**). After SNP calling, we filtered out SNPs with minimum calling rate less than 50% and also filtered out 11/285 accessions because of the high individual missing rate (i.e., > 90%). As a result, a set of 247,008 SNPs were retained for 274 sunflower accessions (see **Supplementary Table S2** and **Supplementary Table S3**). For this SNP set, the average missing rate is 32.3% and the average number of reads per SNP site per sample is 22 (**Supplementary Figure S4**). After filtering the SNP set into 246,671 SNPs by removing the excessive alignments on the 147 unplaced genomic scaffolds, the minor allele frequencies of ≤ 0.05 were applied, resulting in a set of 226,779 SNPs (**Supplementary Table S4**). And these SNPs evenly distribute across the 17 sunflower chromosomes (**Supplementary Figure S5**).

With the filtered SNP set, we found the LD decays rapidly in the population, i.e., average pairwise SNP distance elevated from 30 kb to 18 kb while the LD reduced from *r*^2^ = 0.2 to 0.15 (**Figure 2A**). When the LD is *r*^2^ = 0.1, the average physical distance between two SNPs is about 220 kb, largely consistent with a previous study (58). Principal component analysis suggested that the first principal component (PC) explained about 5% variance and the top 10 PCs explained 25% variance in total (**Supplementary Figure S6**). Using the k-means algorithm, three groups were detected (**Figure 2B**), suggesting our sunflower accessions likely be composed of three sub-populations.

**Figure 2.**
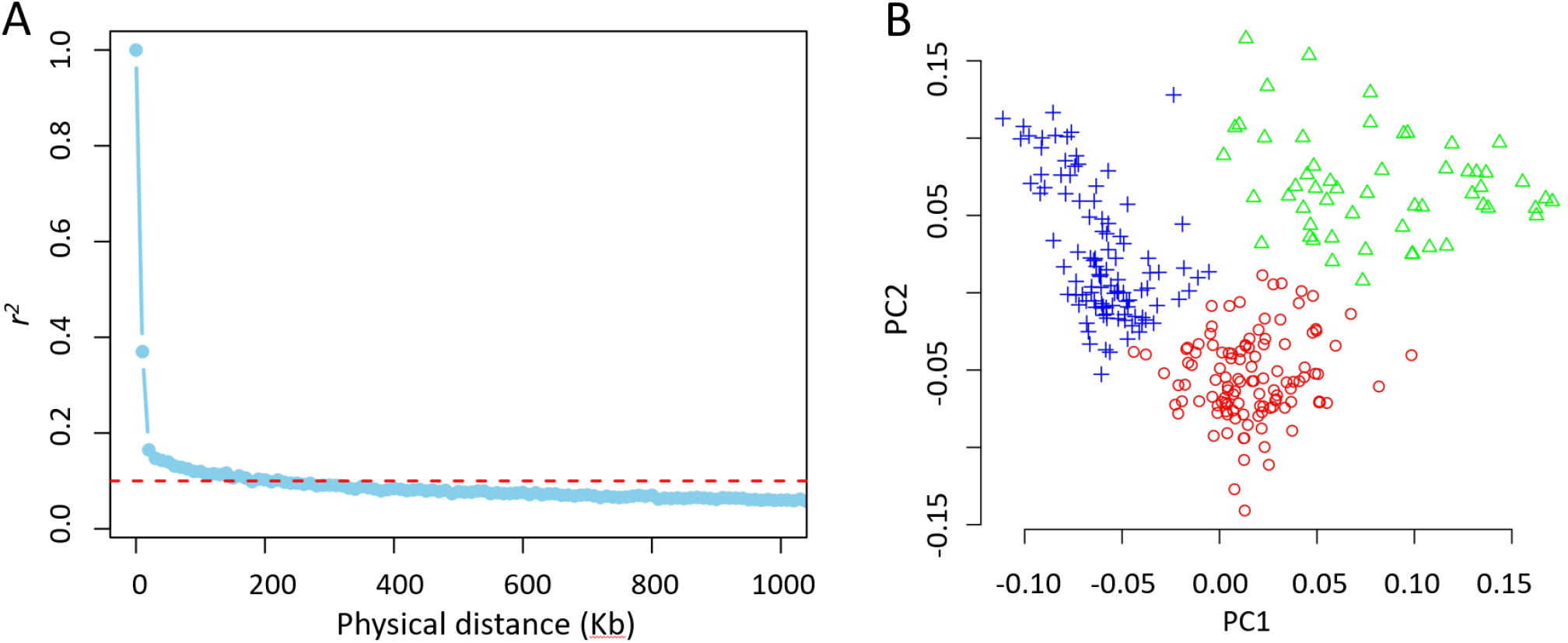
Properties of genotype data. **(A)** LD decay. The threshold was set to 0.1. **(B)** PCA plot. The accessions were clustered by using the simple k-means algorithm with the three groups.

### GWAS results for the head diameter trait

Using the set of 226,779 SNPs, we conducted GWAS with two different statistical methods, i.e., MLM and FarmCPU (see **Materials and Methods**). Quantile-quantile (Q-Q) plots suggested the population structure was well controlled for our GWAS analyses (**Figure 3**). By setting a stringent Bonferroni adjusted P-value cutoff (*P* = 2.2 × 10^−7^ determined by 0.05/n, n = 226,779), FarmCPU method identified *n* = 8 significant trait-associated SNPs located on 6 different chromosomes (**Figure 3A, Supplementary Table S5**). These FarmCPU identified GWAS SNPs were located close to 21 genes (**Supplementary Table S6**. For MLM method, 106 SNPs were identified that were located close to 53 genes (**Supplementary Table S7, Supplementary Table S8**). n=5 GWAS peaks were above the Bonferroni threshold (**Figure 3B**). If requiring at least three significant SNPs within a peak, two significant GWAS peaks were identified, one located on chr10 and the other on chr16. Interestingly, the most significant SNP (NC_035442.2-17467721) on chromosome 10 and another significant SNP (NC_035448.2-31775666) on chromosome 16 found by MLM method were also detected by the FarmCPU method.

**Figure 3.**
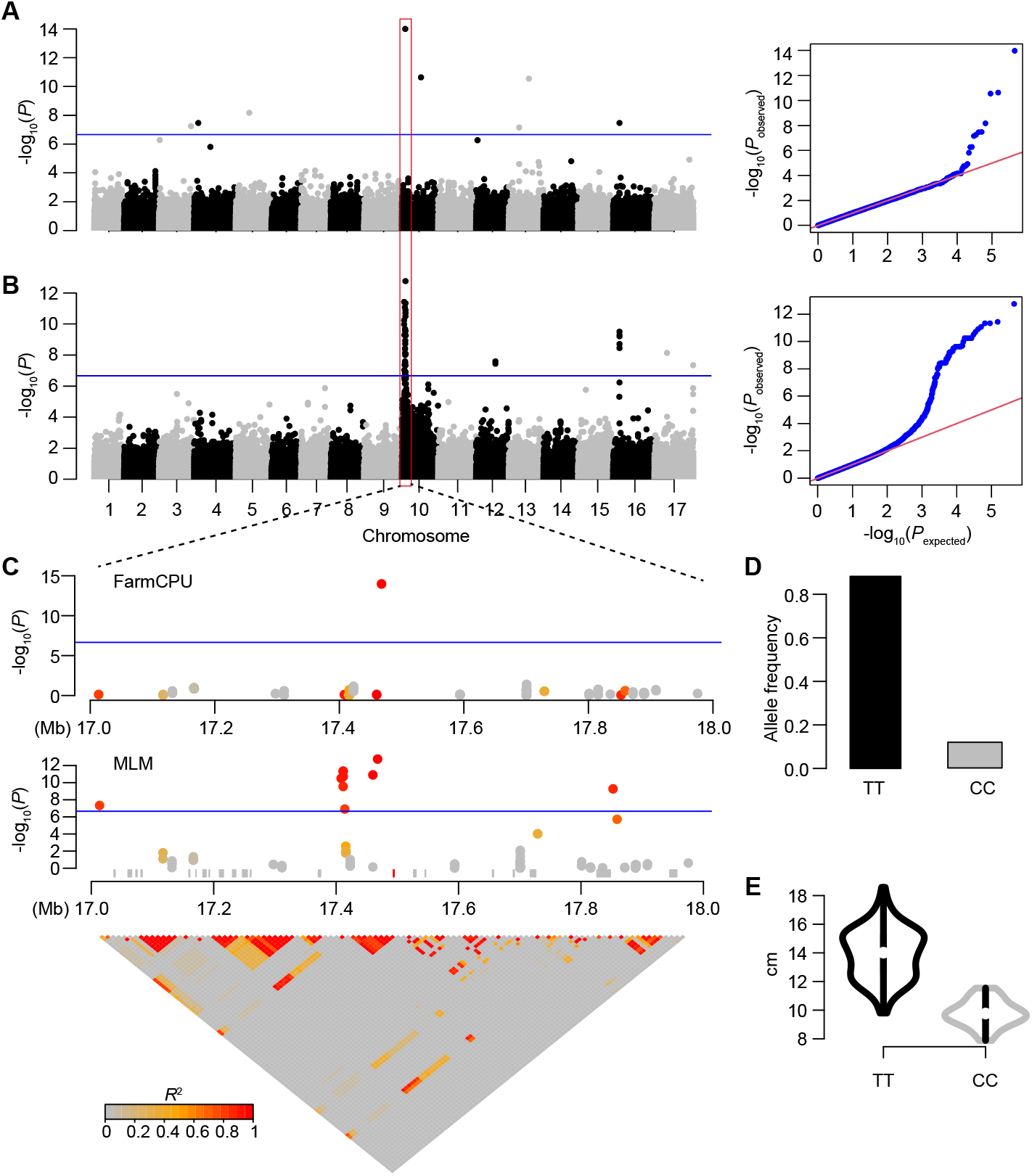
Head diameter associated genetic loci and the zoom-in plots on a Chr10 signal. Manhattan plots and quantile-quantile (Q-Q) plots by using **(A)** FarmCPU and **(B)** MLM methods. The vertical and horizontal axes in the manhattan plots indicate the *P* values in the − *log*_10_ scale and the chromosome numbers, respectively. The horizontal blue lines indicate the significance thresholds. **(C)** A zoom-in plot of the highlighted (500 kb upstream and downstream of the most important SNP) region on chromosome 10 and LD plot in the highlighted region. In the zoo-in plot, the gray rectangles represent the annotated gene models. The red one is the gene that is closest (25kb) to the most significant SNP (NC_035442.2-17467721) at chromosome 10. The LD level (*r*^2^) of windows with the leading signal is represented by color in the zoom-in figure. **(D)** Allele frequency of the most significant SNP detected both in MLM and farmCPU methods. **(E)** Distribution of head diameter trait of sunflower accessions carrying “TT” and “CC” genotypes.

For the chr10 peak, 86 significant trait-associated SNPs were identified, ranging across a 0.9 Mb genomic region from 17.0 Mb to 17.9 Mb (**Figure 3C**). According to the genome annotation information, 41 genes are located within this genomic region. A candidate gene (Gene ID: 118483017) was located 25 kb away from the most significant SNP (NC_035442.2-17467721). The advantage genotype at the most significant GWAS signal is a “TT” genotype, with a frequency of 0.88 (**Figure 3D**). The average head diameter of the sunflower plants carrying the “TT” genotype is 13.93 cm compared to the average value of 9.78 cm for plants with the “CC” genotype (**Figure 3E**).

Similarly, for the peak detected on chromosome 16, 10 significant trait-associated SNPs were identified by using MLM method, one of which was detected by the FarmCPU method (**Figure 3**). The MLM GWAS peak ranges across a 2 Mb genomic region from 31.0 to 33.0 Mb on chromosome 16 and 7 genes are located within this genomic region (**Supplementary Figure S7**). The advantage “TT” genotype at the most significant GWAS signal (NC_035448.2-31753915) has an average head diameter of 13.86 cm with a frequency of 0.91 as compared to the “AA” genotype with the average value of 9.95 cm with a frequency of 0.09.

## Discussion

In this study, we evaluated a set of diverse sunflower accessions for the head diameter trait in replicated field trials over three years. The trait exhibited a large variation and yet was highly heritable with an estimated heritability of 0.88, consistent with previous studies (59). For these diverse accessions, we generated a high-density SNP panel composed of more than 200k high-quality SNPs. As compared to other crop species, the genetic and genomic resources are limited in sunflowers. These diverse sunflower accessions and the high-quality SNP set provide invaluable resources for population genetic analysis in sunflowers.

By a combined analysis of the SNPs and head diameter phenotype, two trait-associated loci were repeatedly detected by using two different GWAS methods. The Chr10 locus can elevate the head diameter by 4 cm, and the Chr16 locus increases by 3 cm. In addition to these two strong GWAS signals, a number of less significant GWAS peaks were identified. Based on these results, markers can be developed to facilitate marker assistant selection. It is anticipated that only by employing the two markers targeting the two major effect loci, a maximum improvement of 7 cm of head diameter can be achieved for a breeding program in which these two loci are dominant by the unfavorable alleles.

In addition to the trait-associated SNPs, we identified a number of genes underlying the GWAS peaks. However, many of these genes have no clear functional assignment due to the limited functional study in sunflowers. To delineate the casual relationship of the trait-associated loci and the head diameter trait variation, further molecular characterization is warranted. As the growing human population and other environmental concerns, sunflower demand has been raising. The GWAS signals and candidate genes can be leveraged to improve the yield and quality of sunflower in order to address the increasing needs.

## Acknowledgements

We thank Dr. Martha Mamo from the Department of Agronomy and Horticulture at University of Nebraska-Lincoln for providing the financial support for partial of the genotyping cost. Yavuz Delen is supported by the Republic of Turkey, Ministry of Education. This work was conducted using the Holland Computing Center of the University of Nebraska-Lincoln, which receives supports from the Nebraska Research Initiative. We thank Musa Ulutas for the helpful discussion.

## Author contributions statement

Y.D.: Data curation, investigation, methodology, visualization, formal analysis, writing - original draft preparation. R.V.M.: Formal analysis, visualization, validation, writing - review & editing. G.X.: Formal analysis, visualization, writing - review & editing. S.P.D.: Investigation, methodology, formal analysis, writing - review & editing. J.C.S.: Writing - review & editing. J.Y.: Supervision, investigation, validation, writing - review & editing. I.D.: Project administration, supervision, conceptualization, investigation, resources, writing - review & editing. All authors reviewed the manuscript.

## Supporting Information

### Supporting Tables

**Table S1. 342 Accessions Phenotyped**. A total of 342 accessions were evaluated in 2019, 2020, and 2022 that were provided by the United States Department of Agriculture, Agricultural Research Center (USDA-ARS), North Central Regional Introduction Station (NCRPIS) in Ames, Iowa, USA (https://github.com/ydelen2/GWAS_Sunflower_Head_Diameter/blob/a4e9a5f230c6ff49ca6ddbc1515eb618c3f88368/Supplementary_Tables/342_accessions_phenotyped.xlsx).

**Table S2. Genotyped accessions by tGBS**. A total of 274 sunflower accessions collected in different regions of the world and stored at the USDA-ARS, NCRPIS were genotyped by tGBS technology (https://github.com/ydelen2/GWAS_Sunflower_Head_Diameter/blob/a4e9a5f230c6ff49ca6ddbc1515eb618c3f88368/Supplementary_Tables/Genotyped_accessions_with_origin.csv).

**Table S3. SNPs detected by tGBS**. After filtering out SNPs with minimum calling rate more than 50% (MCR50), a set of 247,008 SNPs were retained (https://github.com/ydelen2/GWAS_Sunflower_Head_Diameter/blob/766c8692d25b4c054e35153dea57b04d42964052/Data/Sunflower.MCR50.snps.Imputed.hmp.zip).

**Table S4. 226**,**779 SNPs used for GWAS**. After filtering the SNP set into 246,671 SNPs by removing the excessive alignments on the 147 unplaced genomic scaffolds, the minor allele frequencies of ≤ 0.05 were applied, resulting in a set of 226,779 SNPs (https://github.com/ydelen2/GWAS_Sunflower_Head_Diameter/blob/78602b9658f099a7192b4bfa8d78a40e4fc52ad9/Data/Sunflower0.05.recode_226,779_SNPs.zip).

**Table S5. Significant SNPs determined by farmCPU**. The table shows the significant SNPs with position (POS), reference (REF) and alternative (ALT) alleles, SNP effect, standard error (SE), and *p*-value detected by farmCPU method (https://github.com/ydelen2/GWAS_Sunflower_Head_Diameter/blob/a4e9a5f230c6ff49ca6ddbc1515eb618c3f88368/Data/Head_Diameter.FarmCPU_signals.csv).

**Table S6. Gene annotation with significant SNPs detected by farmCPU**. In addition to the detected significant SNPs by farmCPU, it includes the information of genes annotated. Gene annotation was performed using the previously detected genes (http://ftp.ensemblgenomes.org/pub/plants/release-52/gff3/helianthus_annuus/) after applying 50 kb of upstream and downstream to the detected significant SNP positions (https://github.com/ydelen2/GWAS_Sunflower_Head_Diameter/blob/a4e9a5f230c6ff49ca6ddbc1515eb618c3f88368/Data/GWAS_Regions_with_annotation_farmCPU.csv).

**Table S7. Significant SNPs determined by MLM**. The table shows the significant SNPs with position (POS), reference (REF) and alternative (ALT) alleles, SNP effect, standard error (SE), and *p*-value detected by MLM method (https://github.com/ydelen2/GWAS_Sunflower_Head_Diameter/blob/a4e9a5f230c6ff49ca6ddbc1515eb618c3f88368/Data/Head_Diameter.MLM_signals.csv).

**Table S8. Gene annotation with significant SNPs detected by MLM**. In addition to the detected significant SNPs by MLM, it includes the information of genes annotated. Gene annotation was performed using the previously detected genes (http://ftp.ensemblgenomes.org/pub/plants/release-52/gff3/helianthus_annuus/) after applying 50 kb of upstream and downstream to the detected significant SNP positions (https://github.com/ydelen2/GWAS_Sunflower_Head_Diameter/blob/a4e9a5f230c6ff49ca6ddbc1515eb618c3f88368/Data/GWAS_Regions_with_annotation_MLM.csv).

### Supporting Figures

**Figure S1.**
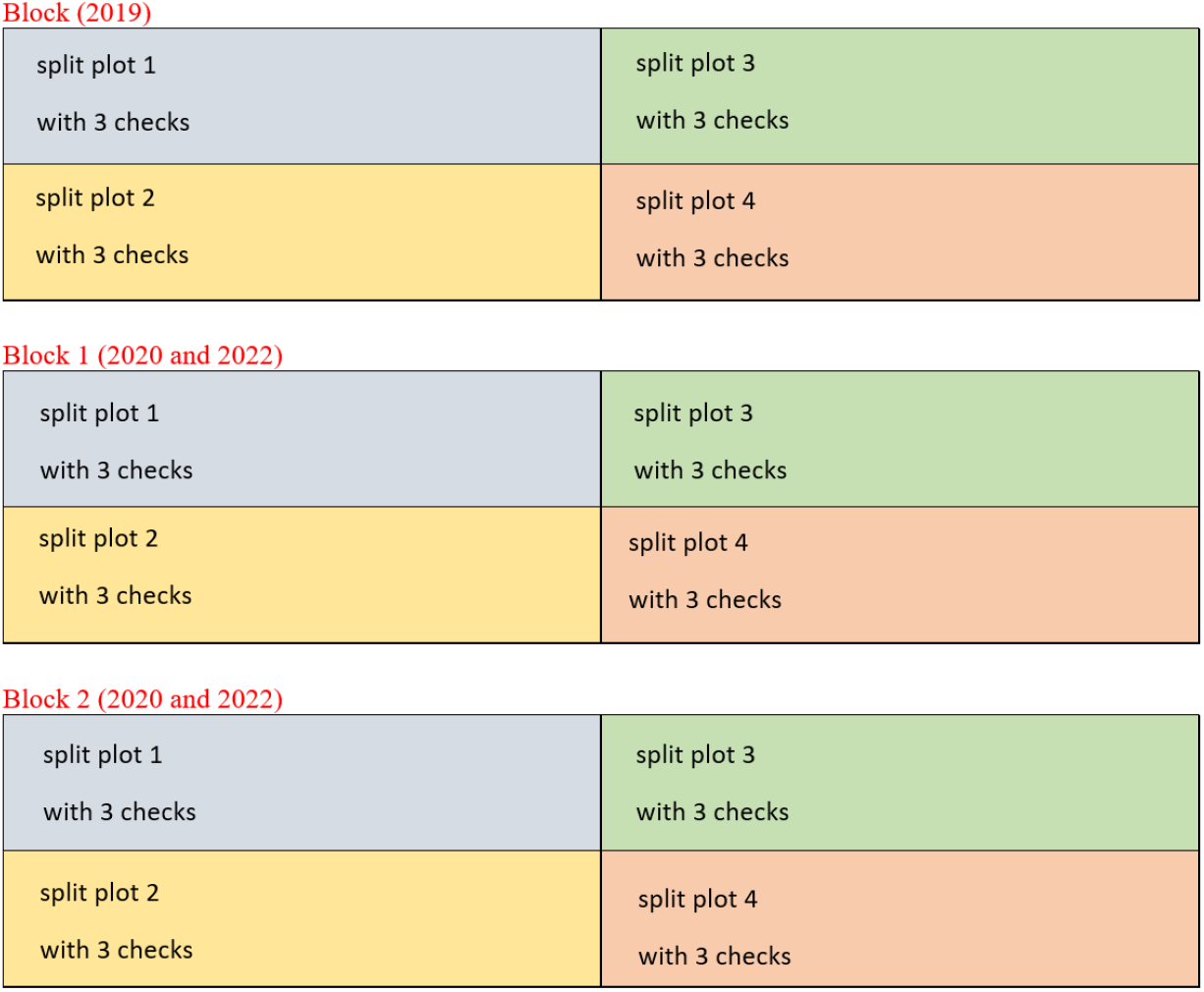
Overview of the field experimental design. An incomplete block design was used with two main blocks (only one block in 2019), four split plots per block, and three replicates of the check (PI 432513) per split plot.

**Figure S2.**
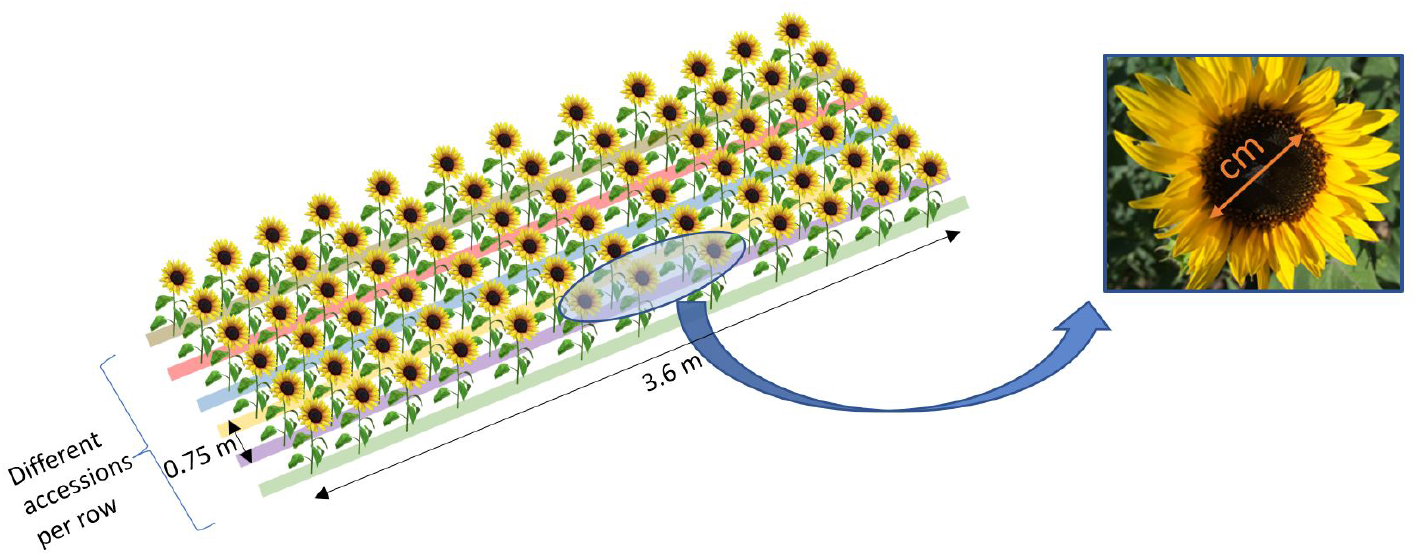
Planting design and phenotype data collection procedure. Head diameter (in centimeters) was measured for three representative plants per plot, excluding edge plants.

**Figure S3.**
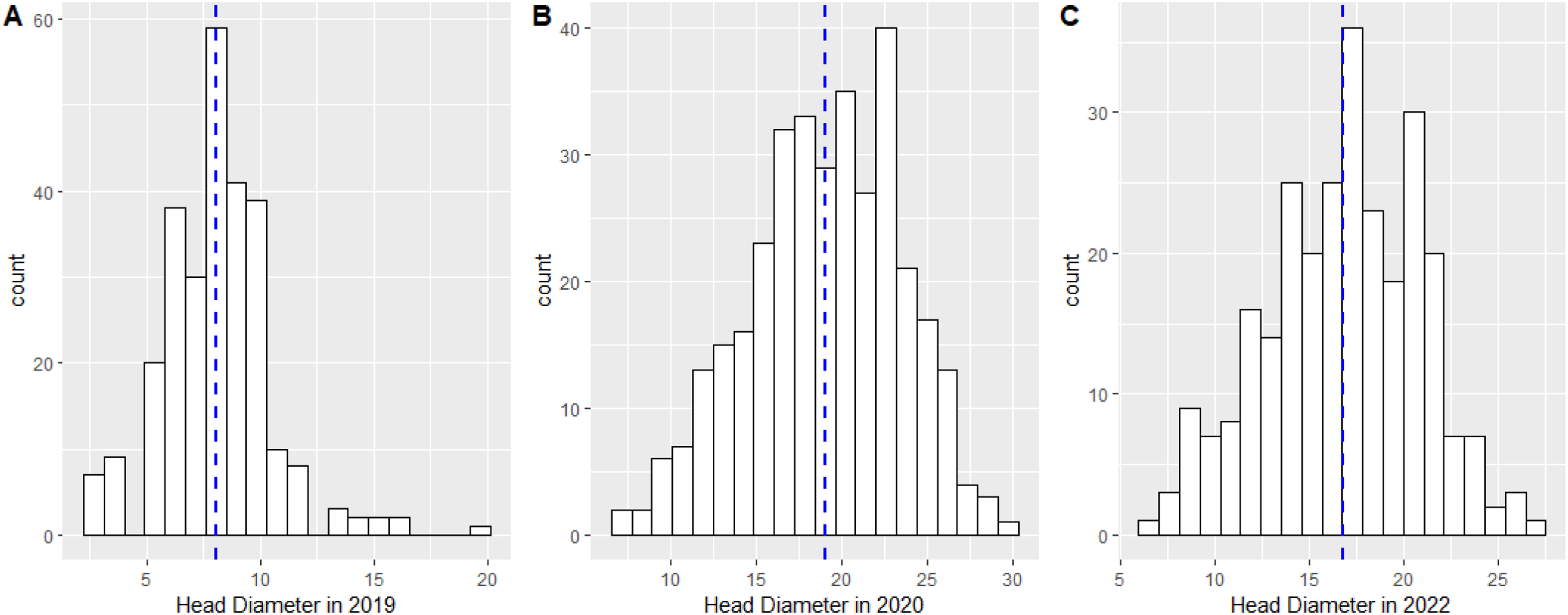
The distribution of average head sizes in 2019 (A) (Average of 3 measurements), 2020 (B) (Average of 6 measurements -2 plots * 3 measurements-), and 2022 (Average of 6 measurements -2 plots * 3 measurements-). The vertical and horizontal axes indicate the number of accessions and the head diameter measurements in cm, respectively. The blue dashed lines show the mean values.

**Figure S4.**
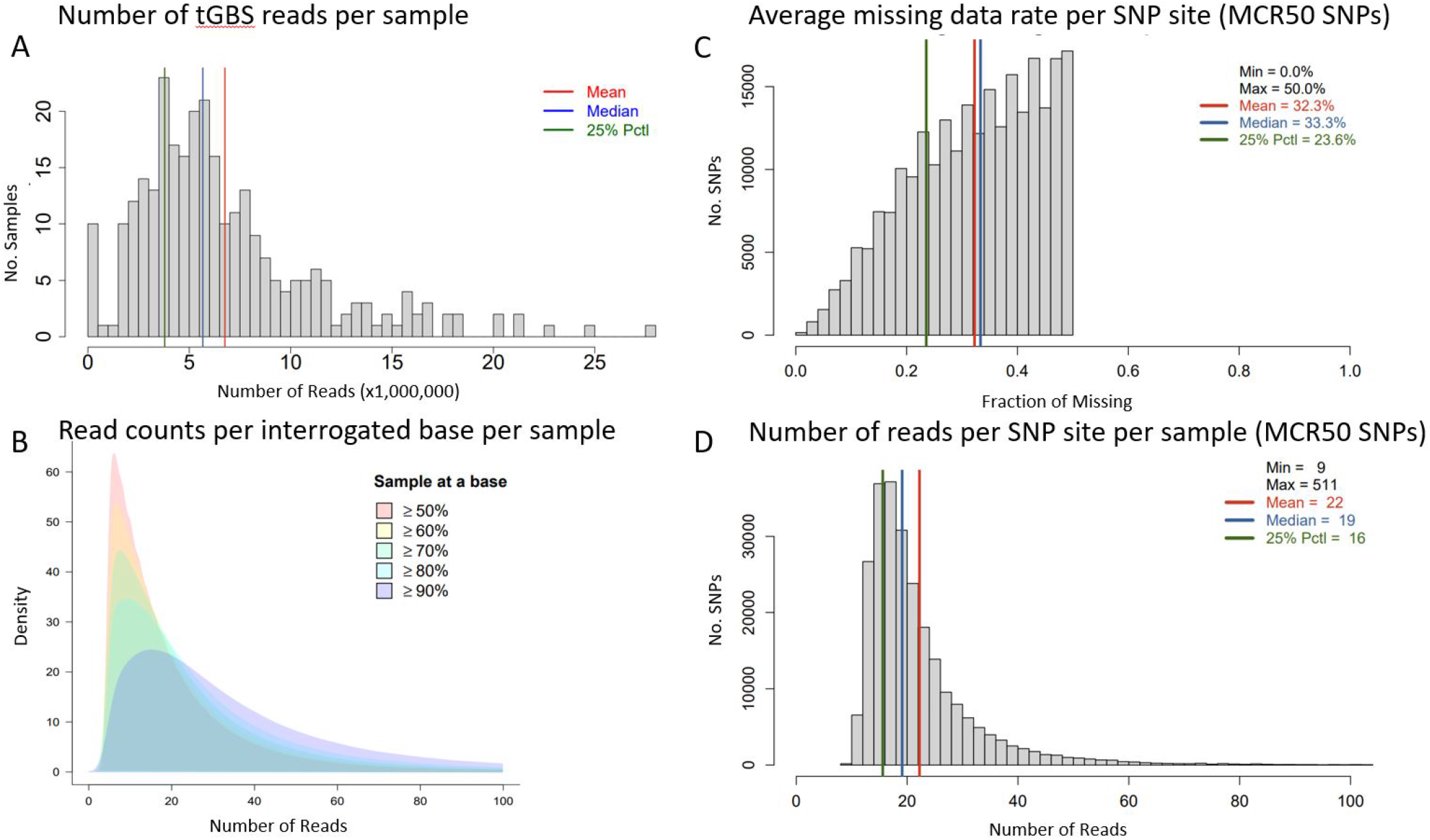
Summary of the genotyping results. **(A)** Distribution of number of reads per sample. **(B)** Distribution of the number of reads per interrogated base per sample. **(C)** Distribution of average missing Rate per SNP site. **(D)** Distribution of the number of reads per SNP site per genotyped sample.

**Figure S5.**
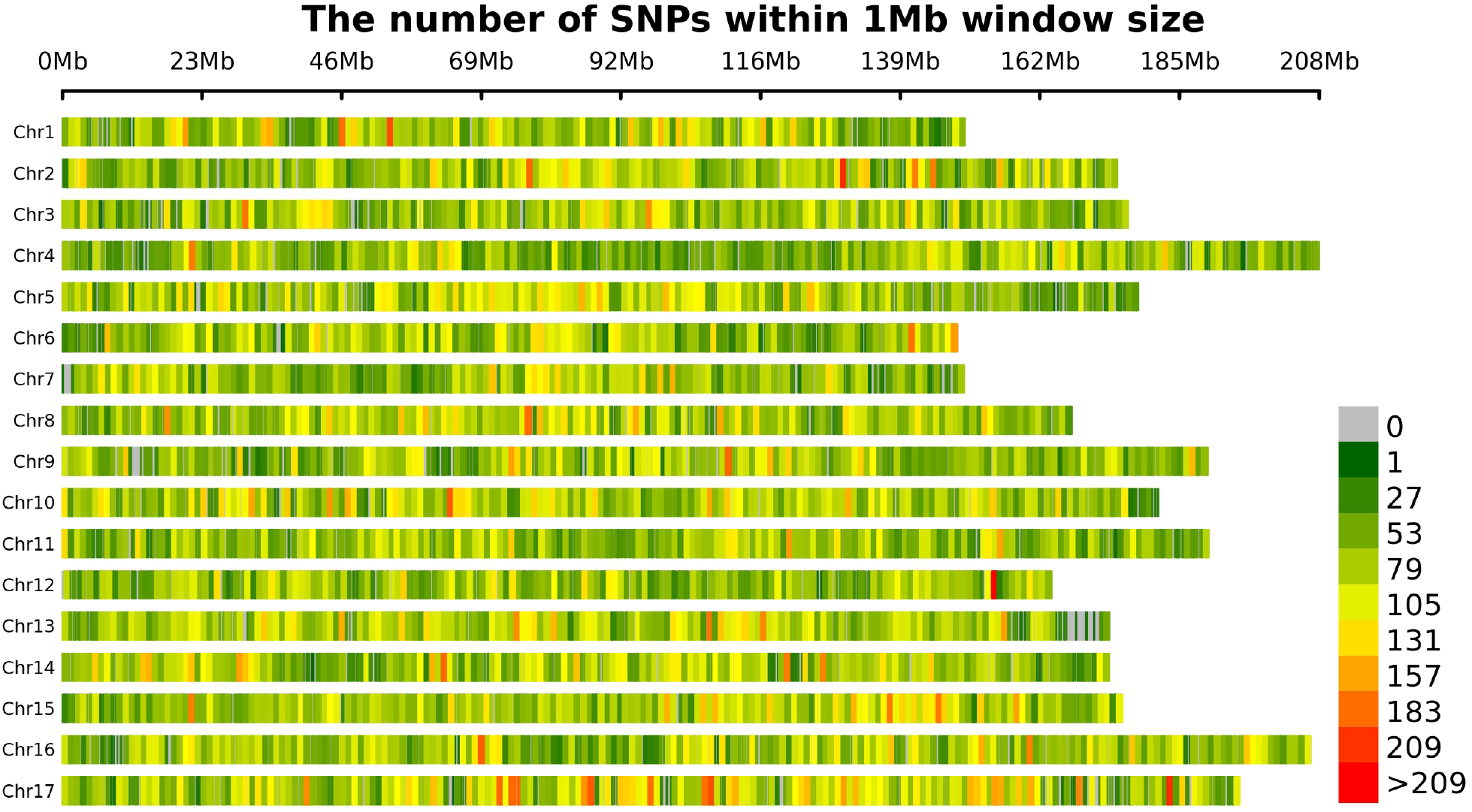
The SNP density distribution on genotyped sunflower population. The number of SNPs within 1 Mb window size through sunflower chromosomes.

**Figure S6.**
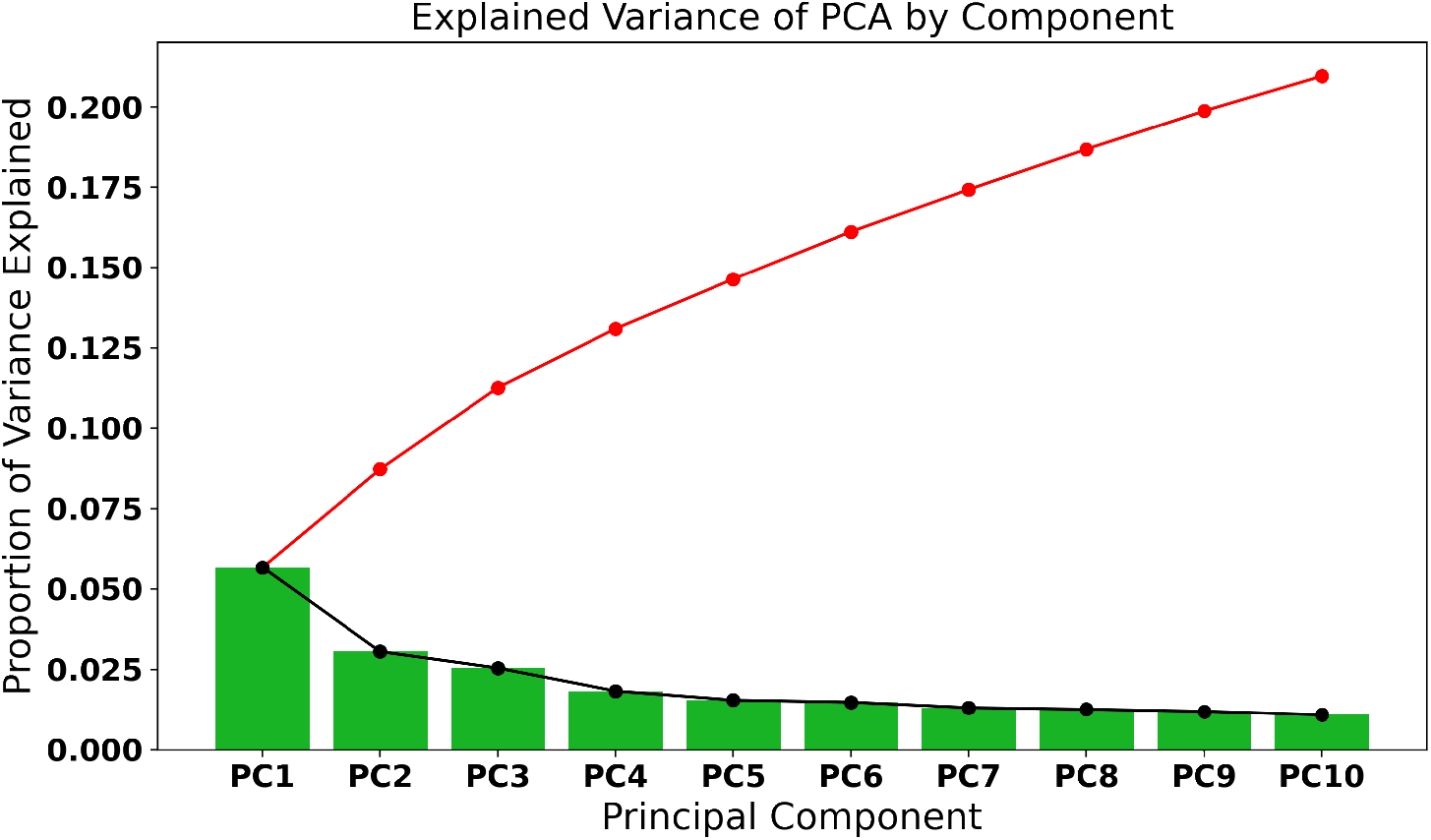
Distribution of explained variance of PCA by component. SNP variation explained by fisrt 10 PCs.

**Figure S7.**
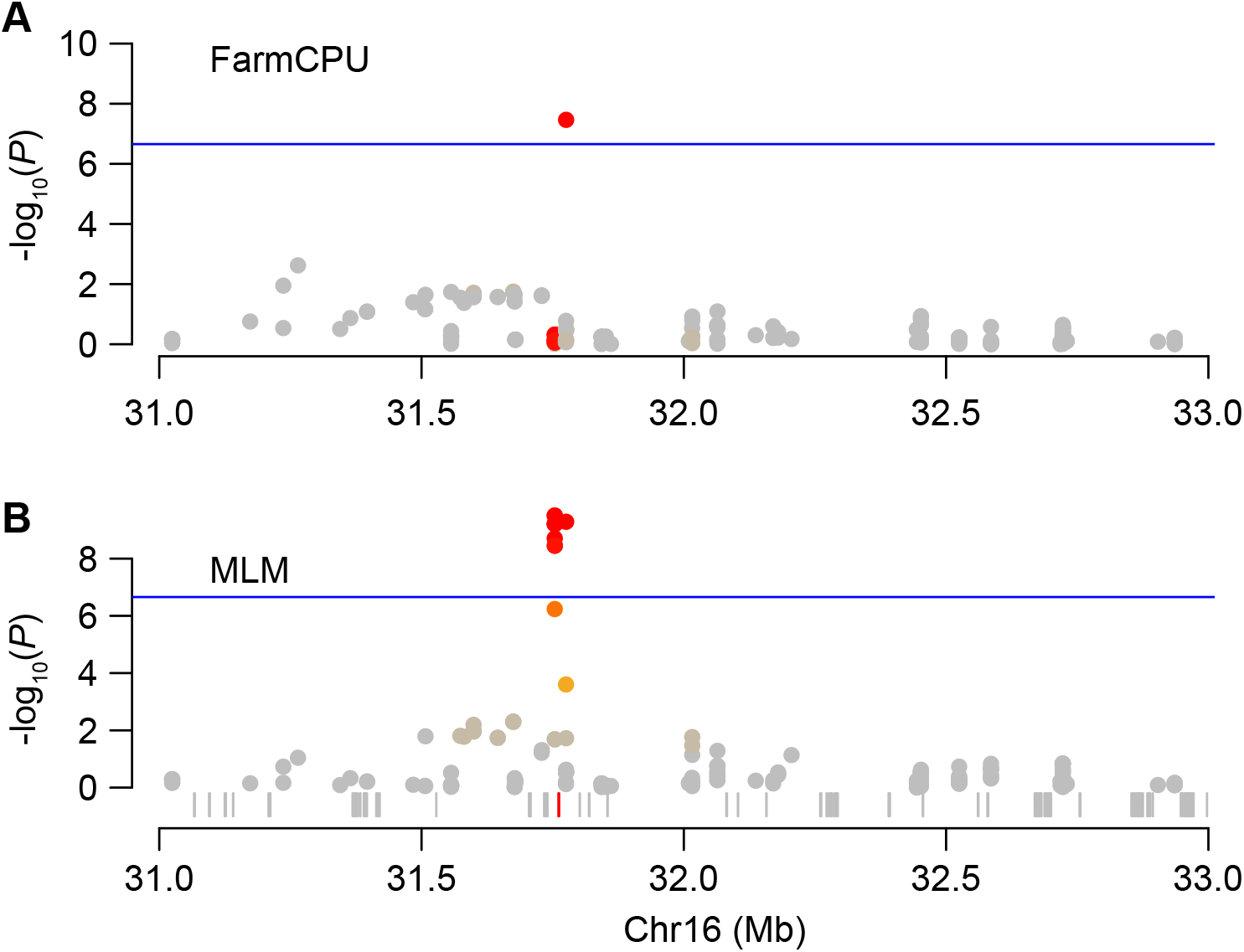
The zoom-in plots of the highlighted (500 kb upstream and downstream of the shared SNP “NC_035448.2-31775666” by farmCPU (A) and MLM (B) methods) region on chromosome 16. The vertical and horizontal axes indicate the *P* values in − *log*_10_ scale and the chromosomal positions, respectively. The points are the SNPs and the blue horizontal line indicates the genome-wide thresholds of Bonferroni correction (*P <* 2.2*e −* 7). The gray rectangles represent the gene models that are annotated. The red one is the gene “LOC110919168” that is closest to the shared SNP at chromosome 16.

